# Isolation and Characterization of 2019-nCoV-like Coronavirus from Malayan Pangolins

**DOI:** 10.1101/2020.02.17.951335

**Authors:** Kangpeng Xiao, Junqiong Zhai, Yaoyu Feng, Niu Zhou, Xu Zhang, Jie-Jian Zou, Na Li, Yaqiong Guo, Xiaobing Li, Xuejuan Shen, Zhipeng Zhang, Fanfan Shu, Wanyi Huang, Yu Li, Ziding Zhang, Rui-Ai Chen, Ya-Jiang Wu, Shi-Ming Peng, Mian Huang, Wei-Jun Xie, Qin-Hui Cai, Fang-Hui Hou, Yahong Liu, Wu Chen, Lihua Xiao, Yongyi Shen

**Affiliations:** College of Veterinary Medicine, South China Agricultural University, Guangzhou 510642, China; Guangdong Laboratory for Lingnan Modern Agriculture, Guangzhou, 510642, China; Guangzhou Zoo, Guangzhou 510070, China; State Key Laboratory of Agrobiotechnology, College of Biological Sciences, China Agricultural University, Beijing, China; Guangdong provincial wildlife rescue center, Guangzhou 510520, China; Zhaoqing Branch Center of Guangdong Laboratory for Lingnan Modern Agricultural Science and Technology, Zhaoqing 526238, China

**Author notes:** Corresponding authors: Yongyi Shen, Lihua Xiao, Wu Chen. These authors contributed equally (Kangpeng Xiao, Junqiong Zhai).

## Abstract

The outbreak of 2019-nCoV in the central Chinese city of Wuhan at the end of 2019 poses unprecedent public health challenges to both China and the rest world^1^. The new coronavirus shares high sequence identity to SARS-CoV and a newly identified bat coronavirus^2^. While bats may be the reservoir host for various coronaviruses, whether 2019-nCoV has other hosts is still ambiguous. In this study, one coronavirus isolated from Malayan pangolins showed 100%, 98.2%, 96.7% and 90.4% amino acid identity with 2019-nCoV in the E, M, N and S genes, respectively. In particular, the receptor-binding domain of the S protein of the Pangolin-CoV is virtually identical to that of 2019-nCoV, with one amino acid difference. Comparison of available genomes suggests 2019-nCoV might have originated from the recombination of a Pangolin-CoV-like virus with a Bat-CoV-RaTG13-like virus. Infected pangolins showed clinical signs and histopathological changes, and the circulating antibodies reacted with the S protein of 2019-nCoV. The isolation of a coronavirus that is highly related to 2019-nCoV in the pangolins suggests that these animals have the potential to act as the intermediate host of 2019-nCoV. The newly identified coronavirus in the most-trafficked mammal could represent a continuous threat to public health if wildlife trade is not effectively controlled.

---

The ongoing COVID-19 outbreak has posed great threat to global public health and caused tremendous economic losses^3^. The virus involved, SARS-CoV-2 (referred in the report subsequently as 2019-nCoV), is one of the three zoonotic coronaviruses (the other two are SARS-CoV and MERS-CoV) infecting the lower respiratory tract and causing severe respiratory syndromes in humans^4,5^. The 2019-nCoV virus was first detected in Wuhan, central China. With the migration of population during the Chinese New Year period, it spreads rapidly from Wuhan to all provinces of China and a few other countries. It has been more contagious but less deadly than SARS-CoV thus far, with the total number of human infections far exceeding that caused by SARS-CoV.

2019-nCoV has about 86.9-88% overall genome sequence similarity to SARS-CoV and bat SARS-related coronaviruses (SARSr-CoV), and clusters with them in phylogenetic analyses of genomic sequences^1,6^. In addition, a bat coronavirus (Bat SARr-CoV RaTG13) with ∼96% sequence identity to 2019-nCoV at the whole-genome level was reported recently^2^. Therefore, it is reasonable to assume that bats are the native host of 2019-nCoV, as previously suggested for SARS-CoVs and MERS-CoVs^7,8^. However, without an intermediate host, bat-derived CoVs rarely infect humans directly. As the outbreak started in December in a big city, hibernating bats were unlikely the direct cause of the epidemic. This raises concerns that whether other hosts of 2019-nCoVs exist and transmitted virus to humans, just like during the SARS and MERS outbreaks.

As coronaviruses are common in mammals and birds^9^, we used the whole genome sequence of 2019-nCoV (WHCV; GenBank accession No. MN908947) in a Blast search of SARS-like CoV sequences in all available mammalian and avian viromic, metagenomic, and transcriptomic data. This led to the identification of 34 highly related contigs in a set of pangolin viral metagenomes (Table S1). Therefore, we have focused our subsequent search of SARSr-CoV in this wild animal.

We obtained the lung tissues from four Chinese pangolins (*Manis pentadactyla*) and 25 Malayan pangolins (*Manis javanica*) that were collected from a wildlife rescue center during March-December 2019, and analyzed for SARSr-CoV using RT-PCR with primers targeting a conservative region of β CoV. RNA from 17 of the 25 Malayan pangolins generated the expected PCR product, while RNA from the Chinese pangolins failed to amplify. In the analysis of plasma samples from eight of the Malayan pangolins using a double-antigen sandwich ELISA designed for the detection of IgG and IgM antibodies against with 2019-nCoV, one sample reacted strongly with an OD_450_ value of 2.12 (cutoff value: 0.11). The plasma remained positive at the dilution of 1:80, suggesting that the pangolin was naturally infected a 2019-nCoV-like virus. Comparing with one β CoV-negative pangolin, histopathological examinations of tissues from four β CoV-positive pangolins revealed the occurrence of interstitial pneumonia in the lung and severe congestion and infiltration of inflammatory cells in the liver, kidney, lymph node, and other organs. Minor hemorrhage was seen in alveolar ducts of the lung and epithelial surface of the bladder (Fig 1 and Fig S1).

**Fig 1.**
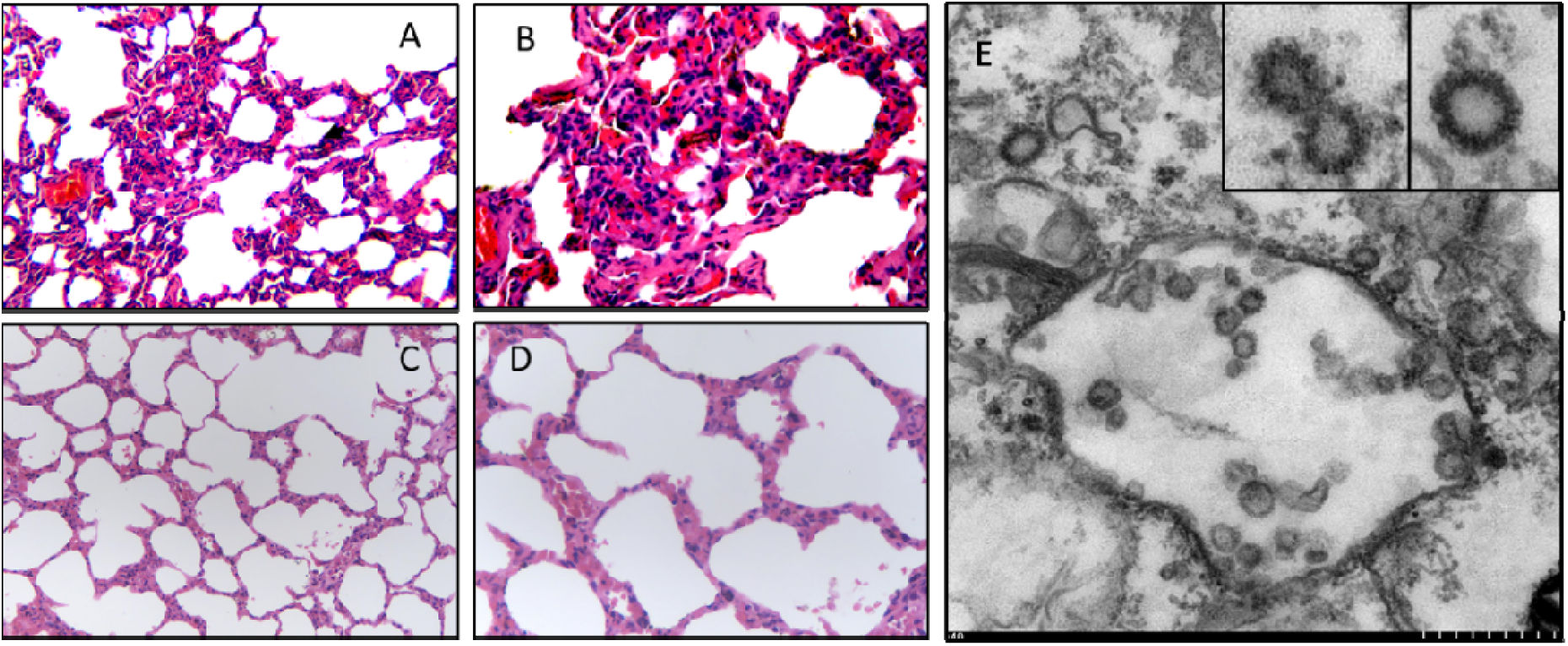
Histopathological (**A-D**) and virological (E) examinations of the lung tissues from an ill Malayan pangolin naturally infected with Pangolin-CoV (**A and B**, 200× and 400×, respectively) and an uninfected Malayan pangolin (**C and D**, 200× and 400×, respectively). Interstitial pneumonia with extensive infiltration of inflammatory cells is seen in the lung tissue from the infected pangolin. Viral particles are seen in double-membrane vesicles in the transmission electron microscopy image (small scale = 50 nm) taken from the Verno E6 cell culture inoculated with supernatant of homogenized lung (**E**), with morphology indicative of coronavirus (inserts at the upper right corner of E).

To isolate the virus, supernatant from homogenized lung tissue from one dead animal was inoculated into Vero E6 cells. Clear cytopathogenic effects were observed in cells after 72 hours incubation. Viral particles were detected by transmission electron microscopy mostly inside double-membrane vesicles, with a few outside them. They showed the typical coronavirus morphology (Fig 1). RT-PCR analysis of the viral culture using five sets of primers targeting the spike (S) and RdRp genes produced the expected PCR products in three of them (Fig S2). The PCR products had ∼84.5% and 92.2% nucleotide sequence identity to the partial S and RdRp genes of 2019-nCoV, respectively (Fig S3).

Illumina RNAseq was used to identify virus in the lung from nine pangolins. The mapping of sequence data to the reference 2019-nCoV (WHCV) genome identified CoV sequence reads in seven samples (Table S2). By *de novo* assembly and targeted PCR, one nearly completed CoV genomes (29,578 Bp) was obtained, and was designated as Pangolin-CoV. The predicted S, E, M and N genes of pangolin-CoV are 3,798, 228, 669 and 1,260 bp in length, respectively, and share 90.4%, 100%, 98.2% and 96.7% amino acid identity to 2019-nCoV (Table 1). In a Simplot analysis of whole genome sequences, the Pangolin-CoV genome was highly similar throughout the genome to 2019-nCoV and Bat SARSr-CoV RaTG13, with sequence identity between 80 and 98%, except for the S gene (Fig. 2). Further comparative analysis of the S gene sequences suggests that there were recombination events among some of the SARSr-CoV analyzed. In the region of nucleotides 1-914, Pangolin-CoV is more similar to Bat SARSr-CoV ZXC21 and Bat SARSr-CoV ZC45, while in remaining part of the gene, Pangolin-CoV is more similar to 2019-nCoV-like and Bat-CoV-RaTG13 (Fig. 2). In particular, the receptor-binding domain (RBD) of the S protein of Pangolin-CoV has only one amino acid difference from that of 2019-nCoV. Overall, these data indicate that 2019-nCoV might have originated from the recombination of a Pangolin-CoV-like virus with a Bat-CoV-RaTG13-like virus (Fig 2). To further support this conclusion, we assessed the evolutionary relationships among β coronaviruses in the full genome, RdRp and S genes, and different regions of the S gene (Fig 3). The topologies mostly showed clustering of Pangolin-CoV with 2019-nCoV and Bat SARSr-CoV RaTG13, with 2019-nCoV and Bat SARSr-CoV RaTG13 forming a subclade within the cluster (Fig 3). In the phylogenetic analysis of RBD region of the S gene, however, Pangolin-CoV and 2019-nCoV grouped together. Conflicts in cluster formation among phylogenetic analyses of different regions of the genome serve as a strong indication of genetic recombination, as previously seen in SARS-CoV and MERS-CoV^5,10^.

**Table 1:** Genomic comparison of Pangolin-CoV with 2019-nCoV, SARS-CoVs and Bat SARSr-CoVs (nt/aa %)

|  | S | E | M | N | Full-length genome |
| --- | --- | --- | --- | --- | --- |
| 2019-nCoV WHCV | 89.1/90.4 | 99.1/100 | 93.8/98.2 | 93.7/96.7 | 90.3 |
| SARS-CoV GD01 | 72.2/77.2 | 93.5/93.5 | 85.8/90.0 | 87.5/90.0 | 81.6 |
| Bat SARSr-CoV RaTG13 | 88.5/89.8 | 99.6/100 | 93.6/99.1 | 94.0/96.7 | 88.9 |
| Bat SARSr-CoV ZC45 | 83.1/86.1 | 98.7/100 | 94.2/99.6 | 88.9/93.3 | 88.0 |
| Bat SARSr-CoV ZXC21 | 81.1/85.4 | 98.7/100 | 94.2/99.6 | 88.9/93.3 | 88.4 |

**Fig 2.**
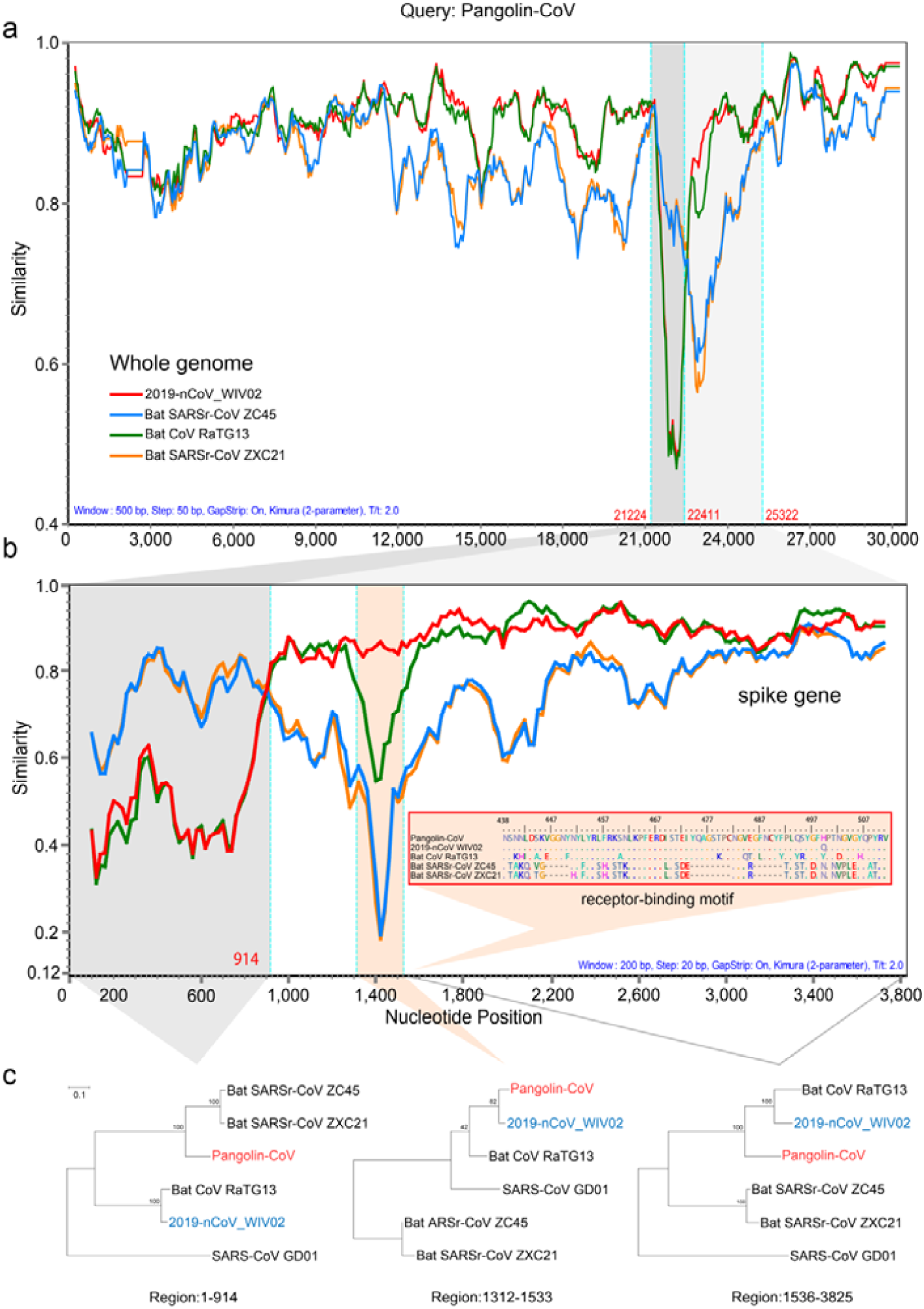
Similarity plot of the near full-length genome sequence (**A**) and S gene sequence (**B**) of Pangolin-CoV to sequences of 2019-nCoV_WIV02, Bat-CoV RaTG13, Bat-CoV ZC45 and Bat-CoV ZXC21. While Pangolin-CoV has a high sequence identity to 2019-nCoV and Bat-CoV-RaTG13 in most regions of the S gene, it is more similar to Bat SARSr-CoV ZXC21 and Bat SARSr-CoV ZC45 at the 5′ end (B). More importantly, Pangolin-CoV is highly similar to 2019-nCoV in the functional RBD region of the S protein, with only one amino difference (insert in **B**). Because of the presence of genetic recombination, there is a discrepancy in cluster formation among the outcomes of phylogenetic analyses of different regions of the S gene (**C**).

**Fig 3.**
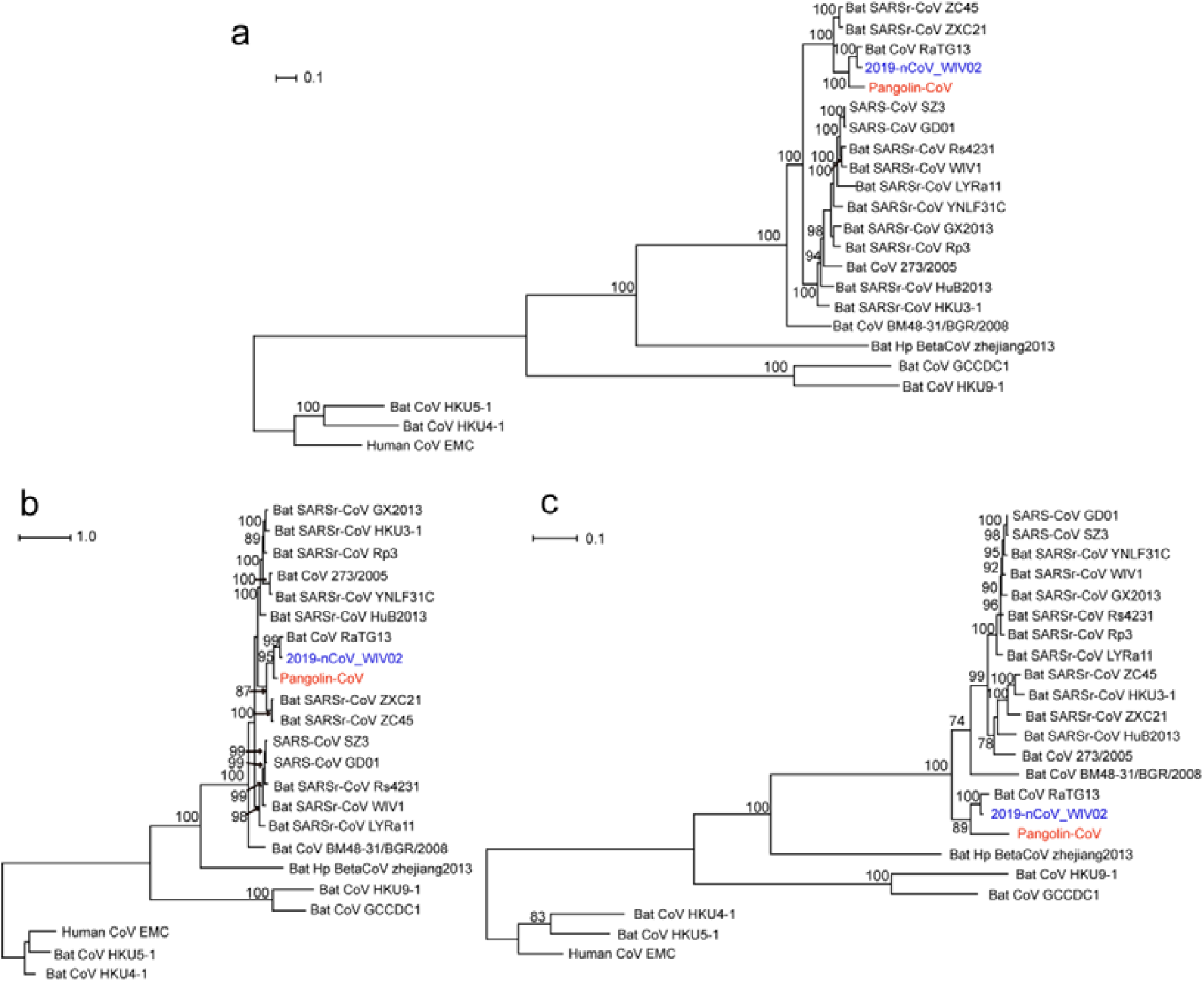
Phylogenetic analysis of nucleotide sequences of coronavirus. (A) Whole genome; (B) Spike gene; (C) RdRp gene.

As the S proteins of both SARS-CoV and 2019-nCoV have been shown to specifically recognize angiotensin converting enzyme II (ACE2) during the entry of host cells^2,11^, we conducted a comparative analysis of the interaction of the S proteins of the four closely related SARSr-CoV viruses with ACE2 proteins from humans, civets and pangolins. As expected, the RBD of the S protein of SARS-CoV bands ACE2 from humans and civets efficiently. In addition, it appears to be capable of binding ACE2 of pangolins. This is also the case for the Bat-CoV-RaTG13. In contrast, the S proteins of 2019-nCoV and Pangolin-CoV can potentially recognize ACE2 of only humans and pangolins. Therefore, ACE2 proteins of humans and pangolins can probably recognize the S proteins of all four SARSr-CoV viruses, while the ACE2 of civets can probably only recognize the S proteins of SARS-CoV and Bat-CoV-RaTG13 (Fig 4).

**Fig 4.**
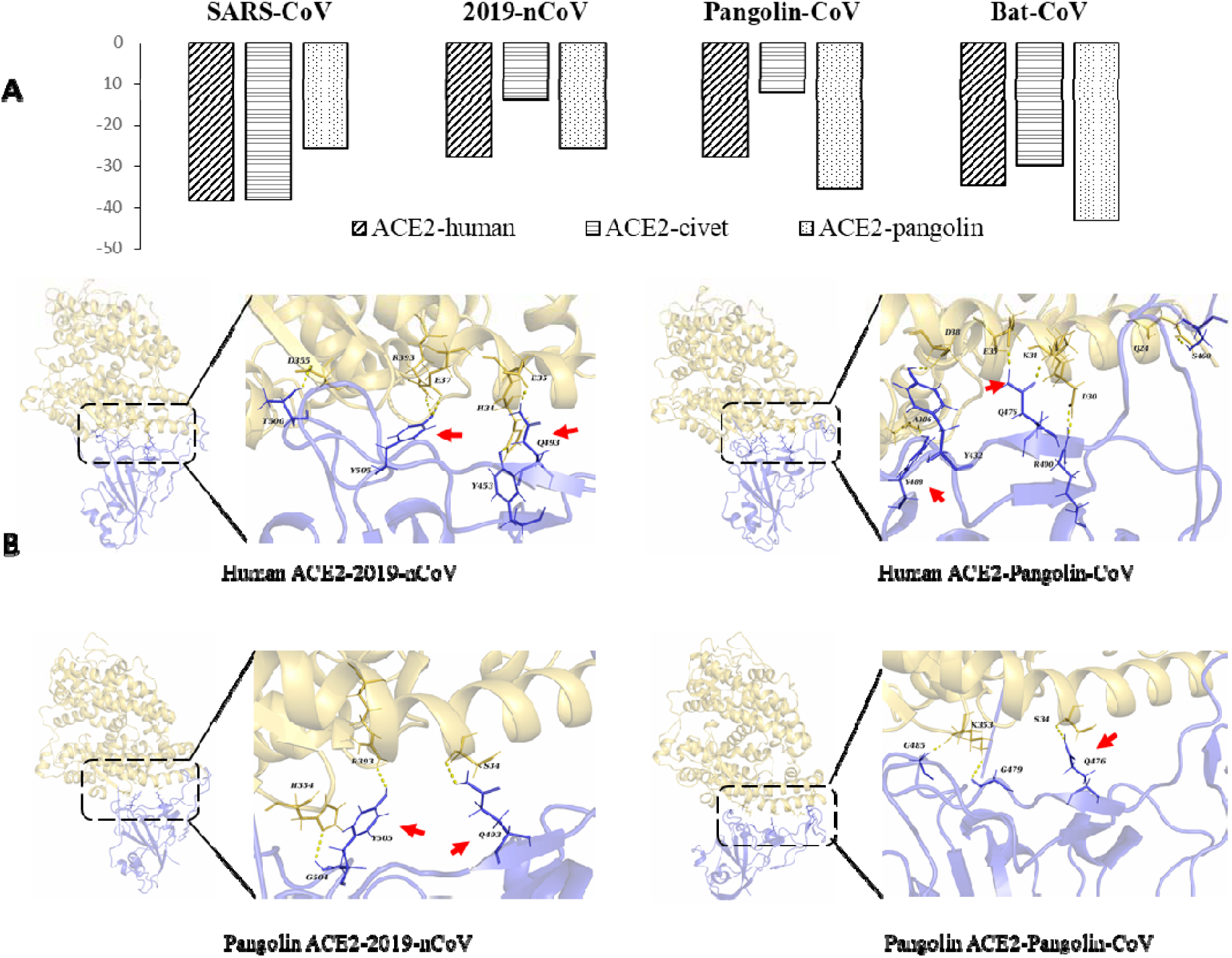
Binding capacity of the RBD of S proteins of four SARSr-CoV viruses to ACE2 of potential hosts. **A.** Free energy (kcal/mol) for the binding between S proteins of four SARSr-CoV viruses and ACE2 of four likely hosts. **B.** Molecular interactions of RBD of the S proteins of 2019-nCoV and Pangolin-CoV with ACE2 proteins of humans and pangolins. The key residues participating in the hydrogen bond formation at the interface between ACE2 and S protein are indicated by red arrows.

Epidemiological investigations of the 2019-nCoV outbreak showed that the initial patients were associated with the Huanan Seafood Market, where live wildlife was also sold^12^. Although the 2019-nCoV virus was detected in some environment samples from the market, no animals thus far have been implicated as carriers of the virus. The SARSr-CoV virus identified in pangolins in the present study is genetically related to 2019-nCoV, but is unlikely directly linked to the outbreak because of the substantial sequence differences between 2019-nCoV and Pangolin-CoV. A virus related to Pangolin-CoV, however, appears to have donated the RBD of the S protein to 2019-nCoV. SARSr-CoV sequences were previously detected in dead Malayan pangolins^13^. These sequences appear to be from Pangolin-CoV identified in the present study judged by their sequence similarity. In the present study, we have provided virological, serological and histopathological evidence for the virulence of Pangolin-CoV and the potential pangolins as the zoonotic reservoir of 2019-nCoV-like coronaviruses. In addition, as the RBD of pangolin-CoV is virtually identical to that of 2019-nCoV, and has strong binding ability to human ACE2, Pangolin-CoV presents a potential future threat to public health. The broad binding range of pangolin ACE2 to RBD of S proteins reaffirms the potential of pangolins as the intermediate host of SARSr-CoV. Pangolins and bats are both nocturnal animals, eat insects, and share overlapping ecological niches, which make pangolins the ideal intermediate host for some SARSr-CoV. Therefore, more systematic and long-term monitoring of other SARSr-CoV in pangolins and other related animals should be implemented to identify the potential animal source of 2019-nCoV in the current outbreak.

Findings in the study support the call for stronger ban of illegal pangolin trade. Due to the insatiable demand for their meat as a delicacy and scales for use in traditional medicine in China, Chinese pangolins are on the verge of extinction (http://www.iucnredlist.org/details/12764/0). The illegal smuggling of other pangolins from Southeast Asia to China is rampant^14,15^. Almost all pangolins in the illegal game market are Malayan pangolins, and the customs confiscates a large number of Malayan pangolins every year. International co-operation and stricter regulations against illegal wildlife trade and consumption of game meat should be implemented. They can offer stronger protection of endangered animals as well as the prevention of major outbreaks caused by SARSr-CoV.

## Materials and methods

### Metagenomic analysis and viral Genome assembly

We collected viromic, metagenomic, and transcriptomic data of different mammals and birds in public databases, including NCBI Sequence Read Archive (SRA) and European Nucleotide Archive (ENA), for searching potential coronavirus sequences. The raw reads from the public databases and some inhouse metagenomic datasets were firstly trimmed using fastp (v0.19.7)^16^ to remove adaptor and low-quality sequences. The clean reads were mapped to the 2019-nCoV reference sequence (MN908947) using BWA-MEM (v0.7.17)^17^ with > 30% matches. The mapped reads were harvested for downstream analysis. Contigs were *de novo* assembled using Megahit (v1.0.3)^18^ and identified as 2019-nCoV-related using BLASTn with E□values < 1e□5 and sequence identity > 90%.

### Samples

Pangolins used in the study were confiscated by Customs and Department of Forestry of Guangdong Province in March-December 2019. They include four Chinese pangolins (*Manis pentadactyla*) and 25 Malayan pangolins (*Manis javanica*). These animals were sent to the wildlife rescue center, and were mostly inactive and sobbing, and eventually died in custody despite exhausting rescue efforts. Tissue samples were taken from the lung, lymph nodes, liver, spleen, muscle, kidney, and other tissues from pangolins that had just died for histopathological and virological examinations.

### Histopathology

Histopathological examinations were performed on tissue samples from five pangolins. Briefly, lung and other tissues collected were cut into small pieces and fixed in 10% buffered formalin for 24 hrs. They were washed free of formalin, dehydrated in ascending grades of ethanol for one hour each, and cleared twice in chloroform for 1½ hours. After being embedded three times with molten paraffin wax at 58°C in a template, the tissue blocks were sectioned with a microtome at 3-μm thickness. The sections were spread on warm water bath (42°C) and transferred onto grease-free glass slides. The slides were air-dried, deparaffinized, and rehydrated through descending grades of ethanol and distilled water for five min each. They were stained with a hematoxylin and eosin staining kit (Baso Diagnostics Inc., Wuhan Servicebio Technology Co., Ltd.) following recommended procedures. Finally, the stained slides were mounted with coverslip and examined under an Olympus BX53 equipped with an Olympus PM-C 35 camera.

### Virus isolation and RT-PCR analysis

Lung tissue extact from pangolins was inoculated into Vero E6 cells for virus isolation. The cell line was tested free of mycoplasma contamination using LookOut Mycoplasma PCR Detection Kit (SIGMA). Cultured cell monolayers were maintained in Dulbecco’s Modified Eagle Medium (DMEM) /Ham’s F-12. The inoculum was prepared by grounding the lung tissue in liquid nitrogen, diluting it 1:2 with DMEM, filtered through a 0.45 μm filter (Merck Millipore), and treated with 16 μg/ml trypsin solution. After incubation at 37□ for 1 hour, the inoculum was removed from the culture and replaced with fresh culture medium. The cells were incubated at 37 □ and observed daily for cytopathic effect.

Viral RNA was extracted from the lung tissue using the QIAamp® Viral RNA Mini kit (Qiagen) following the manufacturer-recommended procedures, and examined for CoV by RT-PCR using a pair of primers (F: 5’-TGGCWTATAGGTTYAATGGYATTGGAG-3’, R: 5’-CCGTCGATTGTGTGWATTTGSACAT-3’) designed to amplify the S gene of β CoV.

### Transmission Electron Microscopy

Cell cultures that showed cytopathic effects were examined for the viral particles using transmission electron microscopy. Cells were harvested from the culture by centrifugation at 1,000× g for 10 min, and fixed initially with 2.5% glutaraldehyde solution at 4 □ for 4 hours, and again with 1% osmium tetroxide. They were dehydrated with graded ethanol and embedded with PON812 resin. Sections (80 nm in thickness) were cut from the resin block and stained with uranyl acetate and lead citrate sequentially. The negative stained grids and ultrathin sections were observed under a HT7800 transmission electron microscope (Hitach).

### Serological test

Plasma samples from eight Malayan pangolins were tested for anti-2019-nCoV antibodies using a double-antigen ELISA kit for the detection of antibodies against 2019-nCoV by Hotgen (Beijing, China), following manufacturer-recommended procedures. In the assay, the coating antigen was S1 protein expressed in eukaryotic cell, and the labeled one was the RBD region of the S protein. The reaction was read on a Synergy HTX Multi-Mode Microplate Reader (BioTek, USA). Positive samples were tested again with serial diluted plasma.

### Metagenomic sequencing

The lung tissue was homogenized by vortexing with silica beads in 1 mL of phosphate-buffered saline. The homogenate was centrifuged at 10,000× g for 5 min and the supernatant was filtered through a 0.45 μm filter (Merck Millipore) to remove large particles. The filtrate or virus culture supernatant was used in RNA extraction with the QIAamp® Viral RNA Mini kit. cDNA was synthesize from the extracted RNA using PrimeScriptScript II reverse transcriptase (Takara) and random primers, and amplified using Klenow Fragment (New England Biolabs). Libraries were prepared with NEBNext^®^ Ultra™ DNA Library Prep Kit for Illumina^®^ (New England Biolabs). Paired-end (150-bp) sequencing was performed on an Illumina NovaSeq 6000. Specific PCR assays were used to fill genome sequence gaps, using primers designed based on sequences flanking the gap.

### Phylogenetic analysis

Multiple sequence alignment of all sequence data was done using MAFFT v7.221^19^. The phylogenetic relationship of the samples was constructed using RAxML v.8.0.14^20^. Best-fit evolutionary models for the sequences in each datasets were identified using ModelTest^21^. Potential recombination events and the location of possible breakpoints in β coronavirus genomes were detected using Simplot (version 3.5.1)^22^ and RDP 4.99^23^.

### Molecular simulation of interactions between RBD and ACE2

The interaction between the RBD of the S protein of SARSr-CoV and ACE2 of humans, civets, and pangolins was examined using molecular dynamic simulation. The crystal structure of SARS-CoV RBD domain binding to human ACE2 protein complex was download from Protein Data Bank (PDB entry: 2AJF^24^). The structures of the complexes formed by ACE2 of civets or pangolins and RBD of 2019-nCoV, Bat-CoV, and Pangolin-CoV constructed using the homology modeling approach on the Swiss-model server^25^, and superimposed with the template (PDB: 2AJF). The sequence identity of SARS-CoV RBD (PDB: 6ACD) to RBD of 2019-nCoV, Bat-CoV and Pangolin-CoV was 76.47%, 76.79% and 74.19%, respectively, while the sequence of ACE2 protein of humans to that of pangolins and civets was 85.40% and 86.93%, respectively.

The molecular dynamic simulations on all RBD-ACE2 complexes were carried out using the AMBER 18 suite^26^ and The ff14SB force field ^27^ as previously described (Liu et al., 2018). After two stage minimization, NVT and NPT-MD, a 10-ns production MD simulation was applied, with the time step being set to 2fs and coordinate trajectories being saved every 1ps. The MM-GBSA^28^ approach was used to calculate the binding free energy of each ACE2 protein to the RBD of CoV S-protein, using the python script MMPBSA.py^29^ in the build-in procedure of AMBER 18 suite. The last 300 frames of all simulations were extracted to calculate the binding free energy that excludes the contributions of disulfide bond. The free energy decomposition method^30^ was used to identify and quantify the key residues that are involved in the interaction between host ACE2 and viral S proteins.

## Acknowledgments

This work was supported by the National Natural Science Foundation of China (Grant No. 31822056 & 31820103014), National Key R&D Program of China (2017YFD0500404), Fund for the Key Program and Creative Research Group of the Department of Education of Guangdong Province, and the 111 Project (D20008), and Ministry of Science and Technology of China, Chinese Academy of Engineering, Chinese Academy of Sciences, Department of Science and Technology of Guangdong Province, and Department of Agriculture of Guangdong Province.

## Author Contributions

YS, LX, and WC conceived the study; JJZ, FHH, YJW, SMP, MH, WJX, QHC and WC collected samples; JZ, NZ, XZ, NL, YG, XL, XS, ZZ, FS and WH performed virus isolation and sequencing; KX, YF, YL, ZiZ, and YS contributed to the analysis; YS and LX wrote the manuscript; YF and RAC edited the manuscript.

## Figure Legends

**Table S1.** Results of Blast search of SARSr-CoV sequences in available mammalian and avian viromic, metagenomic, and transcriptomic data using the 2019-nCoV sequence (GenBank accession No. MN908947)

| Contig name | Sequence identity (%) | length (bp) | e-value | Data source |
| --- | --- | --- | --- | --- |
| con_9 | 98.623 | 363 | 0 | PRJNA573298 |
| con_15 | 97.303 | 519 | 0 | PRJNA573298 |
| con_2 | 96.97 | 203 | 2.41E-44 | PRJNA573298 |
| con_19 | 96.855 | 477 | 0 | PRJNA573298 |
| con_10 | 96.562 | 349 | 4.64E-168 | PRJNA573298 |
| con_1 | 95.96 | 203 | 1.12E-42 | PRJNA573298 |
| con_13 | 95.918 | 373 | 7.65E-42 | PRJNA573298 |
| con_11 | 95.804 | 402 | 5.63E-133 | PRJNA573298 |
| con_4 | 94.737 | 231 | 1.00E-38 | PRJNA573298 |
| con_25 | 94.737 | 779 | 0 | PRJNA573298 |
| con_22 | 94.297 | 655 | 5.73E-115 | PRJNA573298 |
| con_7 | 94.098 | 305 | 1.18E-133 | PRJNA573298 |
| con_16 | 94.041 | 387 | 3.08E-170 | PRJNA573298 |
| con_21 | 93.971 | 680 | 0 | PRJNA573298 |
| con_6 | 93.96 | 304 | 9.13E-130 | PRJNA573298 |
| con_24 | 93.74 | 610 | 0 | PRJNA573298 |
| con_20 | 93.343 | 721 | 0 | PRJNA573298 |
| con_12 | 93.333 | 373 | 4.14E-114 | PRJNA573298 |
| con_30 | 93.321 | 1048 | 0 | PRJNA573298 |
| con_29 | 92.892 | 886 | 0 | PRJNA573298 |
| con_5 | 92.632 | 231 | 2.17E-35 | PRJNA573298 |
| con_31 | 92.495 | 635 | 0 | PRJNA573298 |
| con_23 | 92.354 | 669 | 0 | PRJNA573298 |
| con_32 | 91.942 | 1031 | 0 | PRJNA573298 |
| con_34 | 91.884 | 1687 | 0 | PRJNA573298 |
| con_27 | 91.844 | 846 | 0 | PRJNA573298 |
| con_8 | 91.776 | 304 | 1.99E-121 | PRJNA573298 |
| con_3 | 91.705 | 218 | 6.83E-85 | PRJNA573298 |
| con_14 | 91.436 | 410 | 5.55E-158 | PRJNA573298 |
| con_18 | 91.429 | 385 | 5.23E-153 | PRJNA573298 |
| con_33 | 91.358 | 1177 | 7.06E-27 | PRJNA573298 |
| con_17 | 91.02 | 491 | 0 | PRJNA573298 |
| con_26 | 90.921 | 740 | 0 | PRJNA573298 |
| con_28 | 90.814 | 840 | 0 | PRJNA573298 |

**Table S2.** Identification of SARSr-CoV sequence reads in metagenomes from the lung of pangolins using the 2019-nCoV sequence (GenBank accession No. MN908947) as the reference

| Sample ID | Animal species | Total reads* | No. mapped reads |
| --- | --- | --- | --- |
| M1 | Malayan pangolin | 107,267,359 | 496 |
| M2 | Malayan pangolin | 38,091,846 | 302 |
| M3 | Malayan pangolin | 79,477,358 | 14 |
| M4 | Malayan pangolin | 32,829,850 | 1,100 |
| M5 | Malayan pangolin | 547,302,862 | 56 |
| M6 | Malayan pangolin | 232,433,120 | 10 |
| M8 | Malayan pangolin | 44,440,374 | 12 |
| M10 | Malayan pangolin | 227,801,882 | 0 |
| Z1 | Chinese pangolin | 444,573,526 | 0 |

\*150-bp paired end

## Supplemental Materials

**Fig S1.**
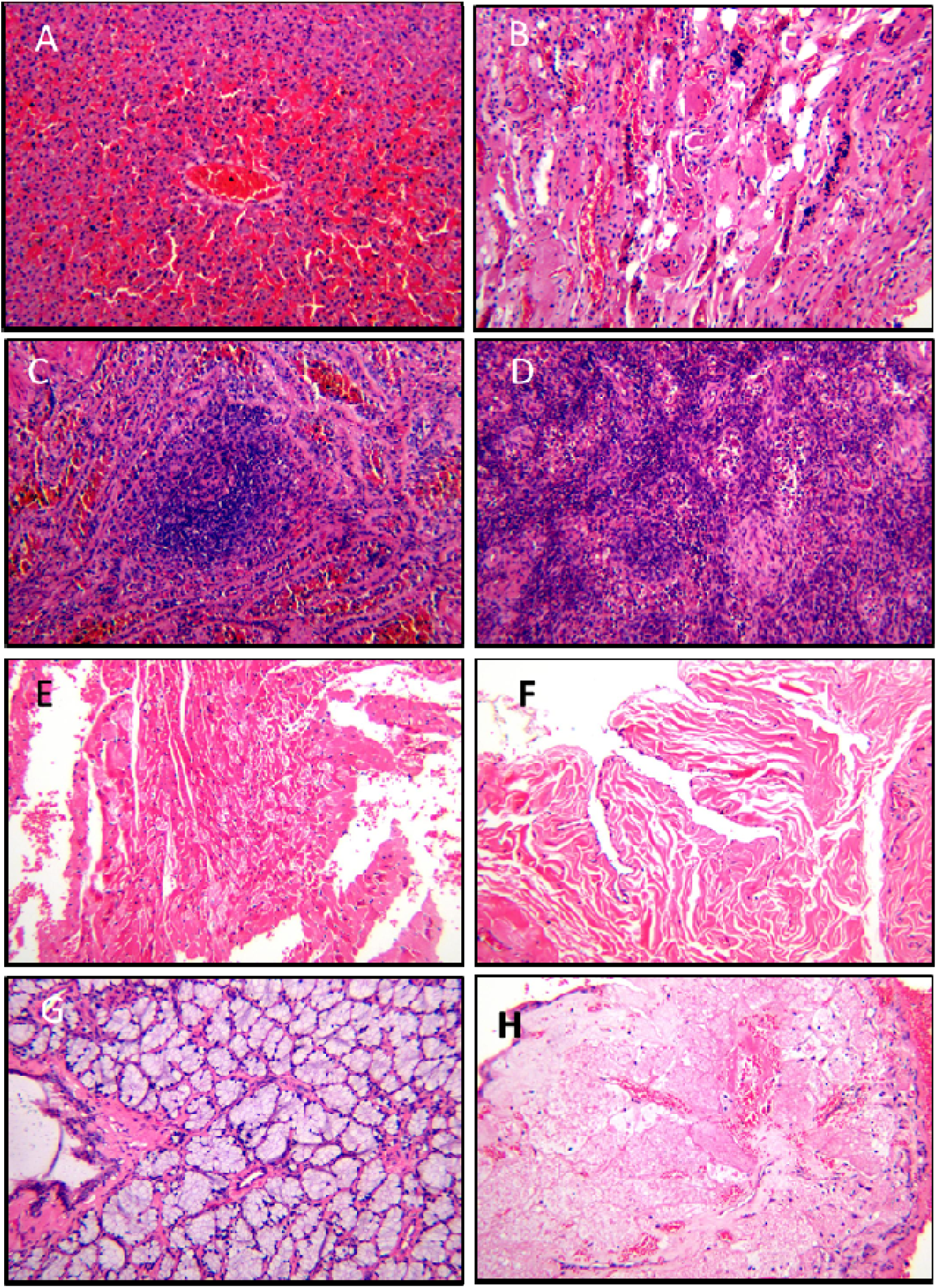
Histopathological changes in liver (A), kidney (B), lymph nodes (C), spleen (D), rectum (E), heart (F), bladder (G), and salivary gland (H) from an ill Malayan pangolin naturally infected with Pangolin-CoV.

**Fig S2.**
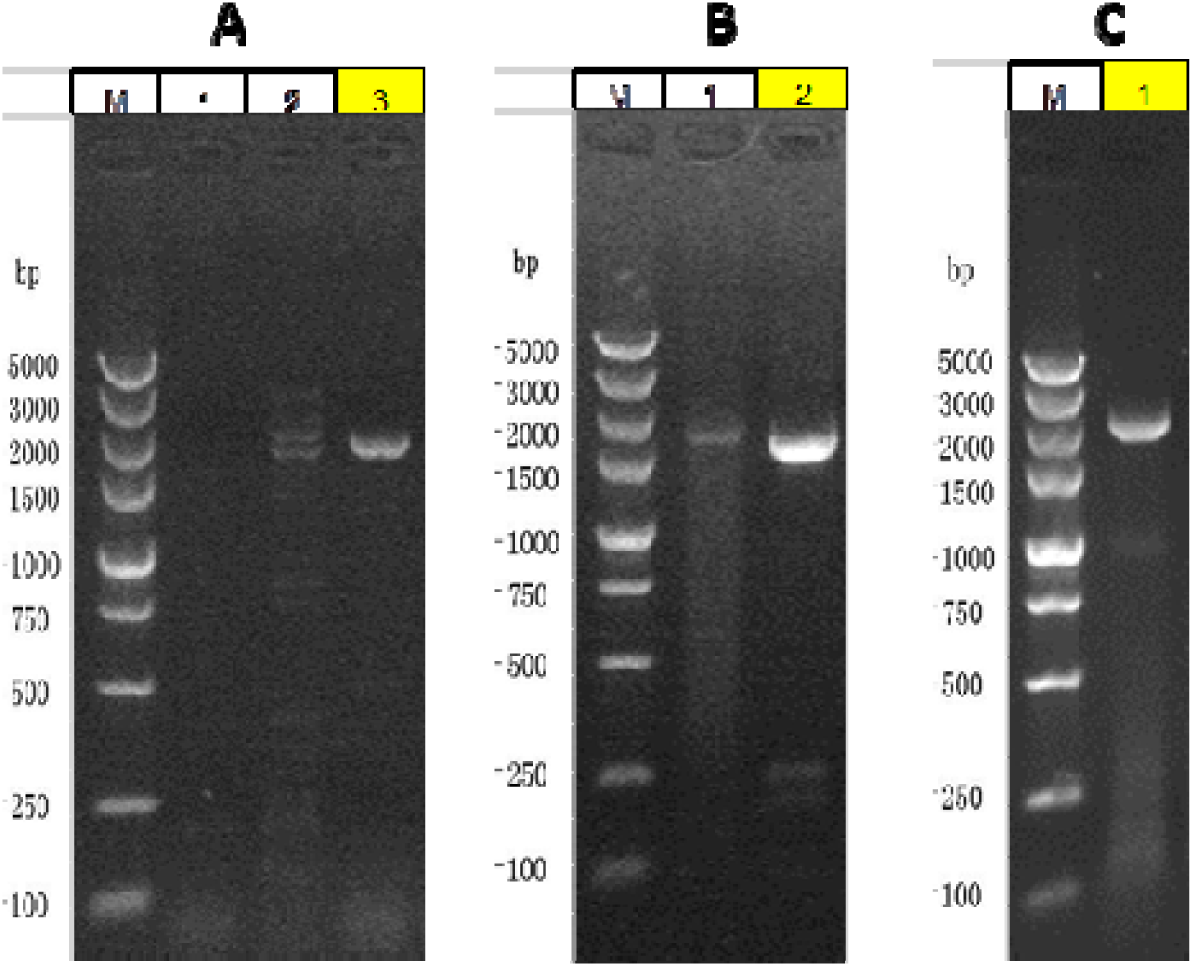
PCR confirmation of the presence of SARSr-CoV in Vero E6 cell culture inoculated with lung tissue of a Malayan pangolin. **(A)** PCR analyses cDNA from viral culture using primers (sets 8, 1 and 6 for lanes 1, 2, and 3, respectively) targeting the S gene. **(B)** Nested PCR analysis (lane 1, primary PCR using primer set 8; lane 2, secondary PCR using primer set 4) of a fragment of the S gene. (C) PCR analysis of the RdRp gene (primer set 10).

**Fig S3.**
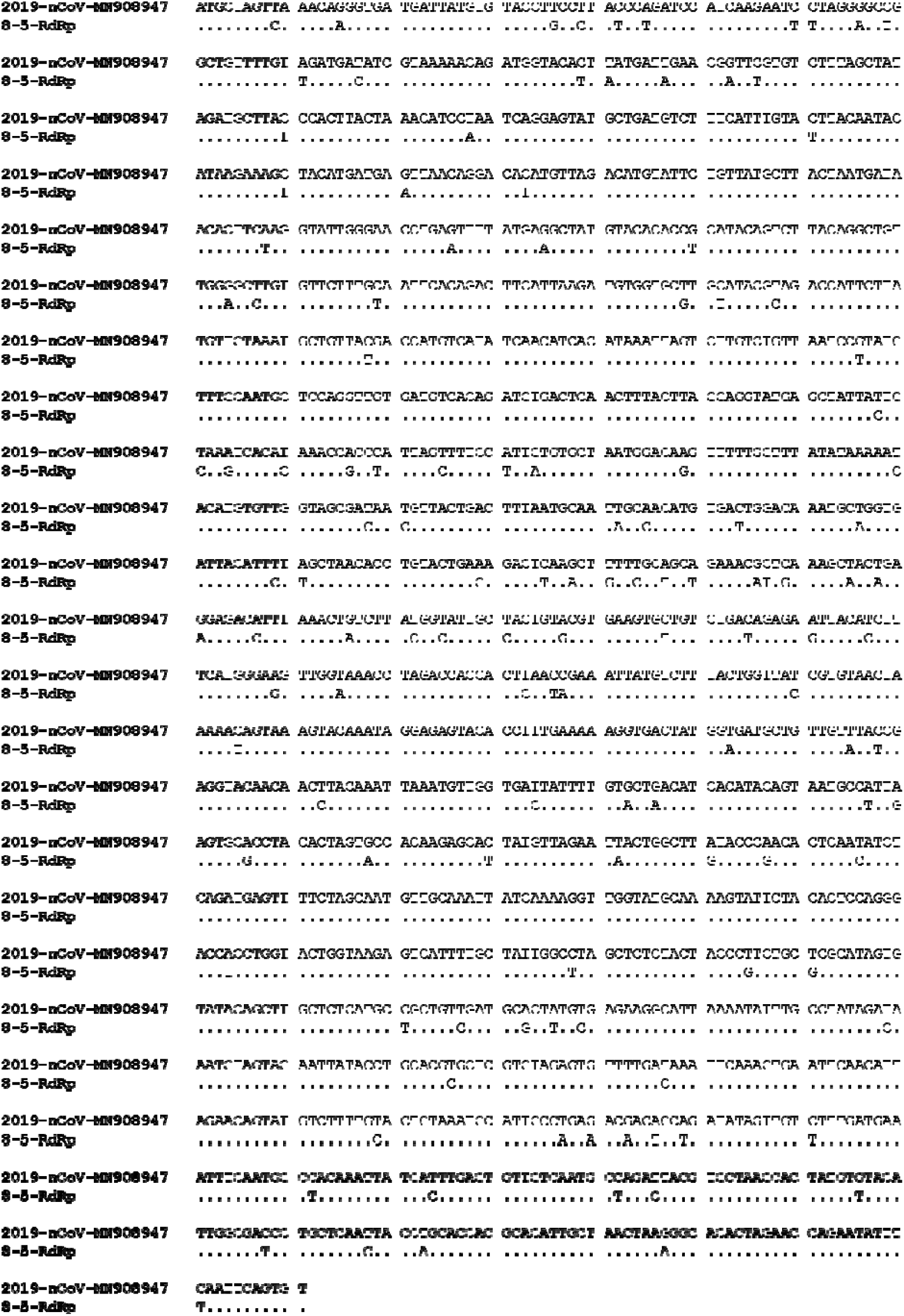
Sequence differences in the partial RdRp gene between 2019-nCoV and Pangolin-CoV. Dots denote nucleotide identity to the reference sequence.

**Fig S4.**
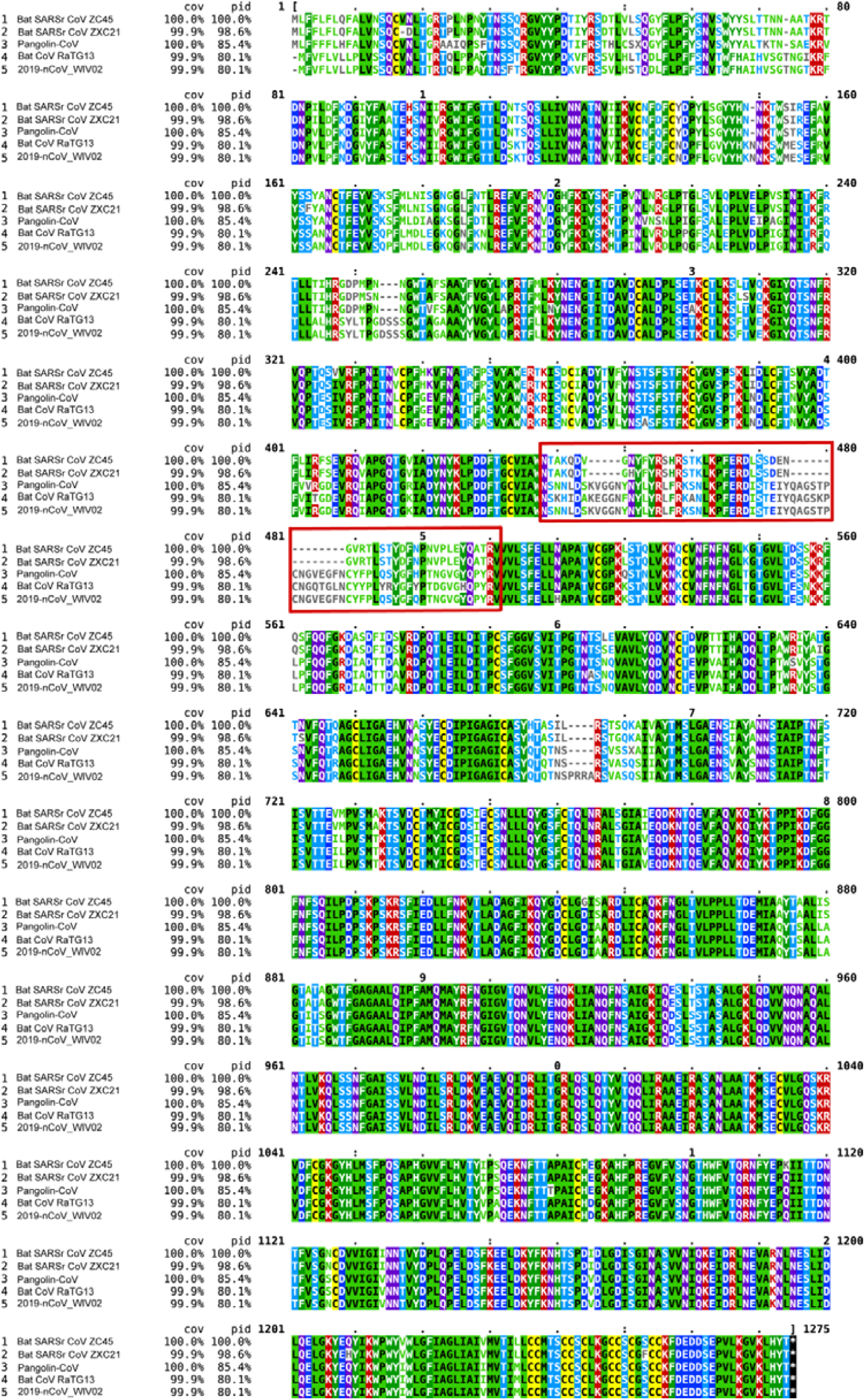
Amino acid sequence alignment of the S protein of the 2019-nCoV with pangolin-CoV, representative bat SARSr-CoVs, and SARS-CoVs.

